# Covariance Nonstationarity is Evident in Spatial Transcriptomics Providing a New Categorization of Spatially Varying Genes

**DOI:** 10.64898/2026.08.10.743911

**Authors:** Puneet Velidi, Zhengxiao Wei, Farouk S. Nathoo

## Abstract

Gaussian process models underlie many spatial transcriptomics tools but typically assume stationary covariance. To test this assumption, we use approximate Bayes factors from R-INLA to compare stationary and non-stationary Matérn covariance functions across 12 Visium datasets; covariance non-stationarity is substantially evident as it appears in 3% to 50% of genes across datasets. We find that gene sets associated with effector functions are enriched among genes favoring non-stationary spatial covariance.

---

Testing for spatially varying genes is an important step in the analysis of spatial transcriptomic data^1^ from both spatial RNA-seq and imaging-based platforms. Spatially varying genes can also be used in downstream analyses that cluster genes based on their estimated spatial processes using algorithms for clustering spatial functional data ^2^. This allows groups of genes that exhibit similar spatial patterns to be identified. These groups are often co-regulated and participate in common biological processes, revealing spatially localized signatures of disease states and, in cancer specimens, distinct tumor microenvironmental niches.

Existing methods primarily model normalized gene expression using Gaussian processes ^3–5^, count data using overdispersed spatial Poisson or negative binomial models ^6–8^, or marks using spatial point-process frameworks ^9,10^. Despite differences in observation models and computational strategies, these methods generally use globally specified spatial dependence structures, often through stationary covariance kernels such as exponential, squared-exponential, Matérn, or periodic covariance functions, and frequently with the additional assumption of isotropy.

Given the heterogeneous nature of many tissue samples and the tendency of local cell populations to vary across the tissue, the stationarity assumption, while statistically and computationally convenient for spatial modeling, may be biologically unrealistic. It is difficult to reconcile this assumption with the pronounced architectural, cellular, and microenvironmental heterogeneity of biological tissues. As a result, biologically meaningful regional changes in spatial dependence may be attenuated or missed, motivating methods that explicitly accommodate non-stationary spatial covariance. More broadly, methods for accommodating non-stationary covariance functions are well developed in spatial statistics ^11–13^, but there has been little application or investigation of non-stationary covariance approaches for spatial transcriptomic data.

In recent work, Sottosanti et al.^14^ consider a Vecchia approximation ^15^ in conjunction with a nearest-neighbor approximation and mixtures of Gaussian processes to cluster spatial transcriptomic data. As far as we are aware, this is the only approach that explicitly accounts for the possibility of non-stationary covariance in spatial transcriptomic data; however, its emphasis is on clustering and point estimation. In contrast, we focus on hypothesis testing, particularly testing for spatially varying genes while accounting for non-stationary spatial covariance.

We test the covariance stationarity assumption across 12 Visium datasets using approximate Bayes factors from R-INLA, comparing Gaussian processes with stationary and non-stationary Matérn covariance functions; genes having expression that is non-stationary are necessarily spatially-varying, albeit according to a covariance function that is more complex than that typically assumed.

All 12 datasets have a non-negligible number of genes that favor non-stationarity (Fig. 1A; Supplemental Material: Table S2). The proportion of genes favoring non-stationarity ranges from 3% to 50%, with 50% of genes in the Human Breast Cancer (ILC) dataset displaying evidence for non-stationarity. Among the genes that favor stationarity, 22% to 93% favor a non-spatial model in the second comparison (Supplemental Data S5). The first comparison therefore shows that the stationarity assumption used by standard tests for spatially varying genes is unwarranted for a substantial and dataset-dependent proportion of genes. The second comparison resembles common practice, except that it is restricted to genes that first favor stationarity, thereby avoiding this form of misspecification.

**Figure 1:**
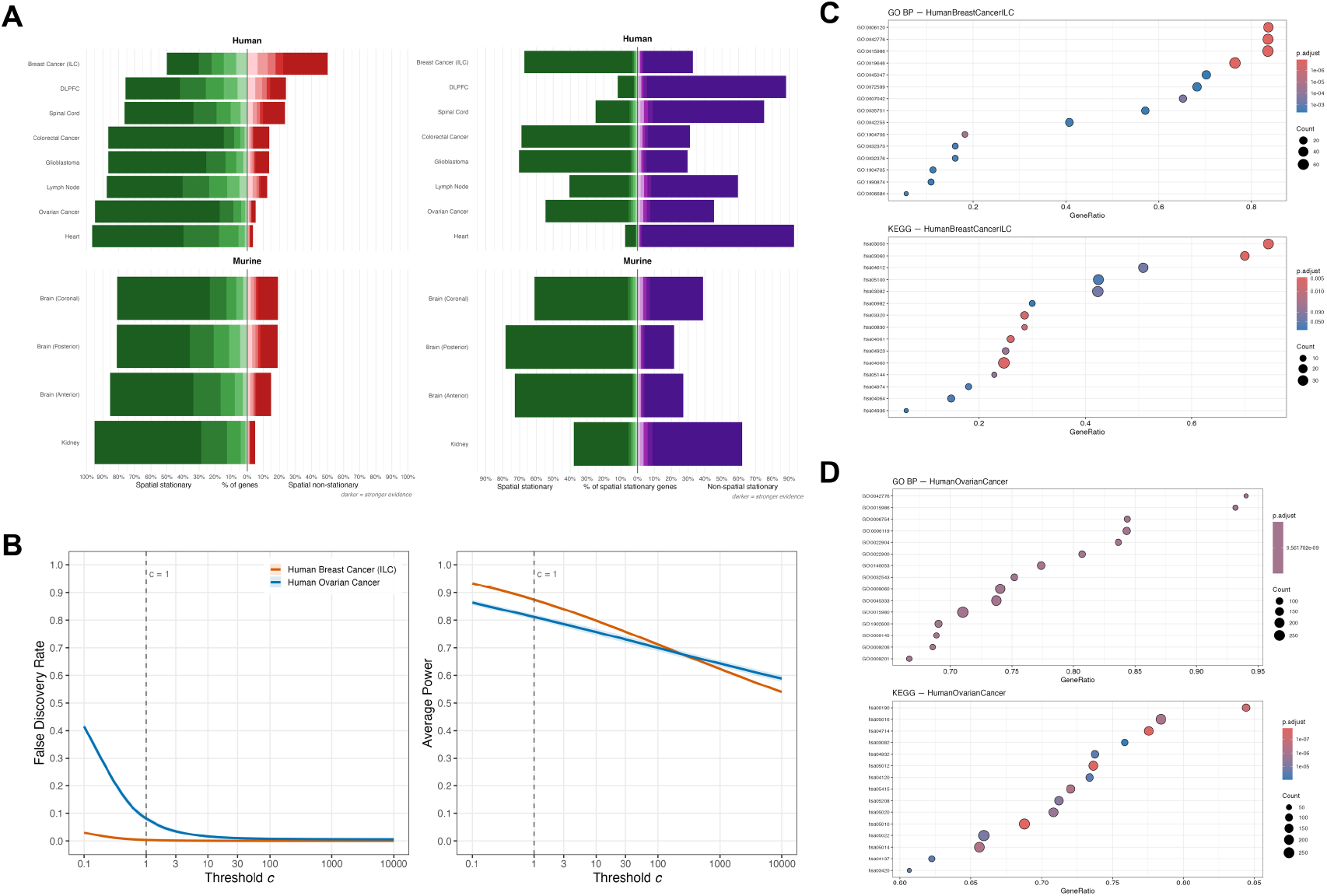
Covariance non-stationarity and gene-set enrichment across 12 Visium datasets. (A) Approximate Bayes-factor model comparisons among successfully fitted genes. The left chart shows the percentage of genes in each dataset favoring stationary or non-stationary spatial covariance. Here, *BF*_10_ = *p*(*y* ∣ *M*_*NS*_)/*p*(*y* ∣ *M*_*S*_), where *M*_*NS*_ is the non-stationary spatial model and *M*_*S*_ is the stationary spatial model; *BF*_01_ = 1/*BF*_10_ favors stationarity. Red stacks indicate genes with *BF*_10_ > 1, and green stacks indicate genes with *BF*_01_ > 1. The right chart is restricted to genes that favored stationarity in the first comparison and shows the percentage favoring the stationary spatial model (green) or the stationary non-spatial model (purple). In both charts, shading indicates evidence strength for the favored model, from lightest to darkest: anecdotal (1 ≤ *BF* < 3), moderate (3 ≤ *BF* < 10), strong (10 ≤ *BF* < 30), very strong (30 ≤ *BF* < 100), and extreme (*BF* ≥ 100). (B) Simulated power and false discovery rate for Human Breast Cancer (ILC) and Human Ovarian Cancer. Curves show the mean of each metric across 100 simulation replicates over Bayes factor threshold *c*, using the decision rule *BF*_10_ > *c* to reject stationarity in favor of non-stationarity. Lighter shaded regions show the mean ± one standard deviation across simulation replicates. (C, D) Gene-set enrichment analysis of genes ranked by *logBF*_10_ for Human Breast Cancer (ILC) and Human Ovarian Cancer, where positive ranks indicate stronger evidence for non-stationary covariance. Enrichment is evaluated using Gene Ontology Biological Process (GO BP) terms and Kyoto Encyclopedia of Genes and Genomes (KEGG) pathways; displayed terms and pathways include enrichment toward both ends of the ranked list. Positive normalized enrichment scores (NES) indicate enrichment toward non-stationarity and negative NES toward stationarity. GeneRatio denotes the number of core-enrichment genes divided by the size of the corresponding tested gene set. Displayed values are Benjamini–Hochberg-adjusted GSEA enrichment-test *p*-values.

We next assess how reliably the Bayes factor procedure distinguishes stationary from non-stationary covariance.

As the Bayes factor threshold becomes more stringent, both the empirical false discovery rate and statistical power decrease, reflecting the increasingly conservative criterion for declaring a gene to exhibit spatially non-stationary covariance. At the chosen threshold of *BF*_10_ > 1, the average statistical power is 0.874 for Human Breast Cancer (ILC) and 0.811 for Human Ovarian Cancer, while the corresponding empirical false discovery rates (FDR) are 0.003 and 0.081, respectively. These datasets represent contrasting prevalences of non-stationary covariance: 50% for Human Breast Cancer and 5.3% for Human Ovarian Cancer. Bayes factors are evidence ratios rather than frequentist *p*-values, and the simulations evaluate the decision rule jointly across all genes thereby demonstrating FDR control under multiple testing. The distributional separation between genes favoring the stationary and non-stationary models in both simulation source datasets is shown in Supplemental Material, Fig. S4. Evidence for non-stationarity has a much wider spread than evidence for stationarity, indicating substantial gene-to-gene heterogeneity: the models are difficult to distinguish for some genes, whereas others provide much stronger evidence for one model.

Having evaluated the testing procedure, we next examine selected residual maps displaying only values below −1 or above +1 (Fig. 2). The non-stationary examples illustrate regions with many threshold-exceeding residuals beside pockets with few displayed values. In the breast-cancer section, *UGT2B11* shows dense positive and negative residuals toward the upper-right, contrasting with a sparse central-left region. In the ovarian-cancer section, *PMP22* shows a subtler contrast between denser positive residuals toward the upper-right and lower areas and fewer displayed values centrally to the left. In the murine anterior-brain section, *Six3* has concentrated positive residuals in the lower region, while most of the upper region has few displayed values. In the murine coronal-brain section, *Dkkl1* shows a curved concentration of positive and negative residuals along the upper-right rim beside a sparse interior. Additional examples use the same fixed residual threshold in Supplemental Material, Figs. S2 and S3.

**Figure 2:**
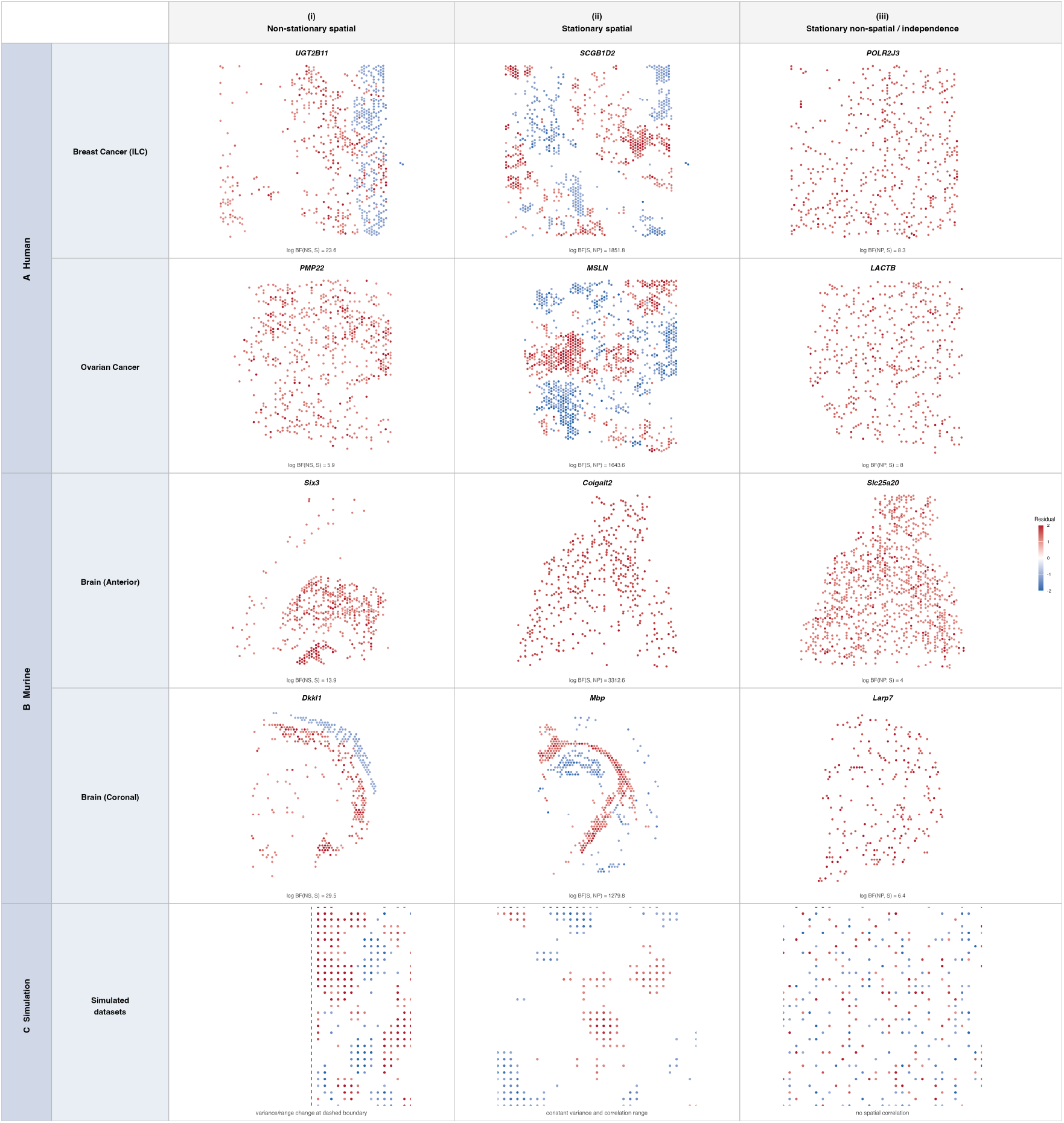
Thresholded residual-expression maps and process simulations. Columns correspond to (i) non-stationary spatial, (ii) stationary spatial, and (iii) stationary non-spatial categories; rows contain (A) two human tissues, (B) two murine tissues, and (C) matched simulated processes, with independence representing the non-spatial category. Observed maps show residuals after quantile normalization and quadratic detrending. Genes were selected for model support and illustrative residual patterns (Methods). Only values below −1 or above +1 are shown. The shared color scale reports residual values (simulated field values in C), centered at zero and clipped to [−2,2]. Values beyond these color limits are retained at the endpoint colors. The dashed line marks the covariance-regime boundary in the non-stationary simulation. S = stationary spatial, NS = non-stationary spatial, NP = stationary non-spatial.

The stationary-spatial genes *SCGB1D2* and *MSLN* show patches of positive and negative residuals, while *Mbp* shows a curved band of positive residuals. The stationary non-spatial examples (*POLR2J3, LACTB, Slc25a20*, and *Larp7*) show dispersed positive residuals at the displayed threshold and were selected for low spatial association in both residuals and squared residuals. All panels use the fixed display threshold |*e*| > 1 and a shared residual color scale clipped to [−2,2]. The simulations in Figure 2C provide known-process comparisons: the non-stationary realization has an empty low-variance region beside clustered positive and negative extremes in the high-variance region; the stationary realization also has clusters, and the independent realization has scattered extremes.

The spatial maps describe individual genes; we next ask whether non-stationarity is also concentrated in coherent biological programs. Figure 1C–D presents the gene-set enrichment results for Human Breast Cancer (ILC) and Human Ovarian Cancer, while the Supplemental Material covers all datasets. Spatial non-stationarity differs strongly across tissues and appears more frequently among genes involved in specialized, context-dependent functions than among broadly expressed housekeeping gene sets. Disease-related pathways are among the strongest enrichment signals, and the leading gene sets in the cancer datasets include processes implicated in disease mechanisms. Among gene sets with positive normalized enrichment scores (NES), immune and cytokine pathways have some of the largest scores across datasets, particularly in heart and colorectal cancer. More generally, these results suggest that gene sets associated with effector functions—those that directly mediate biological responses—are enriched among genes favoring non-stationarity. Neural tissues are enriched for synaptic signaling, neuropeptide activity, gliogenesis, and myelination, while spinal cord tissue shows strong enrichment for ciliary movement, axoneme assembly, and microtubule-based transport. Additional tissue-specific signals include lipid, sterol, and vascular processes in breast cancer and hormone, G-protein-coupled receptor, and epithelial programs in kidney.

Another important consequence of the prevalence of non-stationarity is that a non-negligible set of tissue-segment-specific genes may otherwise be classified as if their spatial variation characterizes the tissue as a whole rather than a particular segment. Tissue segments and their sampling proportions differ among specimens, so conflating segment-specific and tissue-wide signals can reduce cross-sample reproducibility and bias population-level analyses. Non-stationary models do not identify the underlying mechanism on their own, but they can flag genes whose spatial dependence changes locally, providing a new characterization of spatially varying genes. The gene-set enrichment results provide biological context for these genes and show that many belong to tissue-relevant functional programs (Supplemental Material, Table S5).

We note that the prevalence of non-stationarity differs considerably among tissues, suggesting that it may be informative as a tissue characteristic. The Human Breast Cancer (ILC) dataset has the highest prevalence, motivating our closer examination of two human cancer datasets. Cancer can disrupt normal tissue architecture and produce irregular morphology, which may plausibly contribute to locally varying spatial dependence. Future work can investigate the nature of this relationship, but the present analysis of individual tissue sections does not establish a population-level association.

Together, these results suggest that non-stationarity preferentially characterizes tissue-relevant biological programs whose spatial organization varies within the sampled sections. Because gene sets overlap, the data include individual tissue sections, and corresponding histological or cell-type labels are not analyzed here, the enrichments support tissue-specific hypotheses but cannot attribute the observed variation to particular anatomical compartments, cell populations, or cellular mechanisms (Supplemental Material, Tables S5, S6 and Section S2.8).

Covariance non-stationarity, while typically ignored, is evident as a substantial feature in spatial transcriptomic data and provides a new way of characterizing spatially varying genes. Non-stationary spatial approaches allow genes to be categorized as non-spatial, spatially varying according to a stationary covariance, or spatially varying according to a non-stationary covariance.

We anticipate that incorporating tissue anatomy into the covariance structure will further improve both interpretability and detection. These results do not imply that stationary kernels are always inadequate; instead, they show that adequacy is gene- and dataset-specific. Future work will investigate how covariance misspecification affects downstream analyses.

## Methods

### Covariance stationarity and model comparison

A stationary covariance function assumes that the spatial covariance between gene expression measurements *Y*(**s**_*i*_) and *Y*(**s**_*j*_) at spatial locations **s**_*i*_ and **s**_*j*_ depends only on the separation vector **s**_*i*_ − **s**_*j*_. Under covariance stationarity, the strength of spatial correlation can therefore depend on the distance and angle between observation locations, but is otherwise assumed constant across the tissue. Isotropic covariance functions further assume that the spatial covariance depends only on the distance between locations, 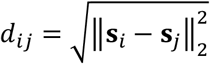.

The consequences of this assumption are especially relevant when testing for spatially varying genes. In the commonly used variance-component formulation, the null model contains only a non-spatial error term, whereas the alternative adds a spatial Gaussian process to the nugget effect. If the true spatial process is covariance non-stationary but the alternative is restricted to a stationary covariance, then the null model itself remains correctly specified when there is no spatial effect, so covariance misspecification is expected primarily to affect power rather than the nominal false-positive rate, provided that the test is appropriately calibrated. Under the alternative, however, the fitted stationary process can only approximate the true spatial dependence, attenuating the likelihood or Bayes-factor evidence separating the spatial model from the non-spatial null and potentially causing genuinely spatial genes to be missed or their spatial scale to be mischaracterized. Spatial statistics has long treated covariance stationarity as an assumption to be assessed rather than an innocuous default, and has developed both flexible non-stationary covariance models and formal tests of stationarity for this reason ^12,16,17^. This distinction has received comparatively little attention in spatial transcriptomics, where increasingly sophisticated observation models are still commonly combined with globally stationary representations of spatial dependence.

### Comparison with conditional autoregressive models

INLAomics ^18^ is a recently developed conditionally Poisson mixed model for spatially varying protein and gene expression data, in which spatial effects are modeled using a multivariate conditional autoregressive (MCAR) prior. While computationally convenient, the conditional autoregressive (CAR) model can lead to counterintuitive spatial dependence structures, particularly when applied to data collected at irregular spatial locations ^19^. It is unclear whether these spatial dependence structures are appropriate or interpretable for gene expression. This criticism of the CAR model applies to the irregularly shaped tissue samples depicted here (Supplemental Material: Fig. S1) and to spatial transcriptomic data more generally.

### Non-stationary Bayesian testing for spatially varying genes

A geostatistical non-stationary Matérn framework underlying the spatial model allows for a more straightforward and intuitive interpretation of the spatial covariance structure. The standard isotropic Matérn covariance structure is a special case, allowing us to test for non-stationarity of the spatial covariance using the Bayes factor (Methods; Supplemental Methods, Sections S2.3–S2.6).

The model for normalized gene expression is a linear model with a spatial effect assumed to follow a Gaussian process with a non-stationary Matérn kernel. Models of non-stationarity can be either isotropic (direction-agnostic) or anisotropic (direction-dependent); for this paper, we focus on the simpler isotropic case. A useful and computationally scalable approach to modeling non-stationary spatial processes is the stochastic partial differential equation (SPDE) representation of the Matérn process ^20^. Its approximate solution uses Gaussian Markov random fields, with approximate Bayesian inference based on integrated nested Laplace approximations (INLA) ^21^. In the stationary and isotropic case, the covariance function takes the form

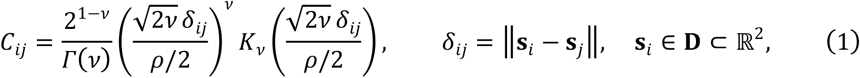

where *K*_*v*_ is the modified Bessel function of the second kind of order *v* > 0. Under the reparameterization 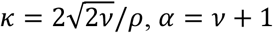, and 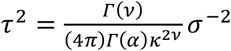, a non-stationary process arises by allowing *κ* and *τ* to vary across location **s**,

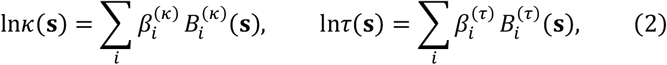

where {*B*_*i*_(**s**)} is a set of spatial basis functions. This specification is well established, and approximate Bayesian inference for the process is implemented in standard software. Further model specifications and implementation excerpts are provided in Supplemental Material, Sections S2.4 and S2.5.

The model is fitted within the integrated nested Laplace approximation (INLA) framework. Two additional models—one with a simpler stationary Whittle–Matérn field and another without a spatial effect—are also considered. The marginal likelihoods associated with each of the three models are estimated using INLA ^22^ and combined with prior model probabilities to produce posterior probabilities for non-stationary spatial variation, stationary spatial variation, and no spatial variation for a given gene.

### Sequential model comparisons

After gene expression normalization and quadratic spatial mean-detrending (Supplemental Material, Section S2.2), we first test spatial non-stationarity versus stationarity. For genes that appear to have stationary spatial covariance, we further test whether they are spatially varying at all by computing the Bayes factor relative to a non-spatial model (Supplemental Methods, Section S2.6).

Testing non-stationarity versus stationarity, followed by stationarity versus non-spatial can be viewed as model selection moving from the most complicated form of spatial covariance towards the simplest (non-spatial) form. Moving in the opposite direction is not recommended because in that case the first test is a comparison of two misspecified covariance models for genes with expression variability corresponding to a non-stationary covariance function.

### Simulation design and calibration

A Bayes factor measures how much more strongly the observed data support one model than another (Supplemental Material, Section S2.6). To evaluate the operating characteristics of the R-INLA Bayes factor test, we simulate expression data for every successfully fitted, simulation-eligible gene in two datasets. We use the fitted non-stationary model when *BF*_10_ > 1 and the fitted stationary model otherwise, then reapply the testing procedure to the simulated data. This allows us to estimate empirically the statistical power and false discovery rate associated with the Bayes factor decision threshold (Supplemental Material, Section S2.7) when parameters are set according to estimates from real data.

In Figure 1B, we simulate 100 replicate gene expression datasets for each gene using the model and parameter estimates selected by INLA from the original Human Breast Cancer (ILC) and Human Ovarian Cancer datasets. We then refit both competing models to each simulated dataset and estimate the empirical false discovery rate and statistical power across a range of Bayes factor decision thresholds (Supplemental Material, Section S2.7 and Table S4). Throughout the manuscript, we adopt a decision threshold of *BF*_10_ > 1, indicated by the dotted grey line in Figure 1B.

### Residual-map examples and illustrative simulations

For the observed examples in Figure 2, non-stationary genes were selected among those with *BF*_NS,S_ > 10 for contrasts in local residual variance and the density of threshold-exceeding residuals. Non-spatial examples required *BF*_S,NS_ > 10 and *BF*_NP,S_ > 10 and were selected for low spatial association of residuals and squared residuals. Stationary-spatial examples required *BF*_S,NS_ > 10 and *BF*_S,NP_ > 1. These descriptive, post-selection criteria identify illustrations, not independent tests of the corresponding spatial properties; screening details and fixed gene selections are documented in the Supplemental Material and analysis code.

For the illustrative thresholded maps in Figure 2C, we simulated one realization on a 31 × 31 regular grid from each of three zero-mean Gaussian processes: a non-stationary Matérn process with an abrupt change in marginal variance and correlation scale at the vertical midline, a stationary Matérn process with constant unit variance and correlation scale, and an independent standard Gaussian process. For both observed residual maps and simulated-process maps, only values below −1 or above +1 are displayed. Colors represent actual residual or simulated field values on a shared diverging scale centered at zero and clipped to [−2,2]. Observations outside the color limits are retained, with their colors saturated at the corresponding endpoint. The fixed display threshold changes only the visualization; model fitting uses all analyzed in-tissue spots. Full covariance specifications and reproducibility settings are provided in Supplemental Material, Section S2.2.

### Data and preprocessing

The complete analysis contained 196,612 gene–dataset fits; 102 optimizer failures (0.052%) were retained as explicit missing-result rows and excluded from model-category percentages.

We analyze 12 publicly available human and mouse 10x Genomics Visium tissue sections distributed as SpatialExperiment objects through ExperimentHub and associated Bioconductor resources. Source count matrices have been processed using Space Ranger; dataset-specific versions and accession information are reported in Supplemental Material, Table S2. Duplicate re-releases are excluded, including MouseBrainCoronal in favor of the Visium_mouseCoronal representation of the same section. Spots explicitly marked in_tissue == 1 are retained; objects without an in_tissue field are treated as already tissue-restricted and retain all supplied spots. This selection retains 3,639 of 4,992 spots for Visium_humanDLPFC and 2,702 of 4,992 spots for Visium_mouseCoronal. Genes are retained if they are detected in at least 0.5% of analyzed tissue spots and have a total count of at least three across the section. For each gene, counts are transformed using log1p and mapped to normal scores using a rank-based inverse-normal transformation. A quadratic trend in the centered spatial coordinates is then removed, and the resulting residuals are used for the spatial covariance analyses. Optimizer failures are retained as explicit missing-result rows; summaries and category percentages use successfully fitted genes as their denominator. Complete dataset and preprocessing details are provided in Supplemental Material, Sections S2.1 and S2.2; dataset-specific mesh settings are reported in Supplemental Material, Table S3.

## Ethics statement

This study involved secondary analysis of publicly available, de-identified human spatial transcriptomic datasets and publicly available mouse datasets. No new participants were recruited and no new human specimens were collected. Ethics approvals and consent procedures for the original studies are described in the corresponding source records and publications. Separate ethics approval was not required for the present secondary analysis.

## Use of generative artificial intelligence

ChatGPT (OpenAI) was used to assist in identifying potentially relevant literature and copyediting (formatting, proofreading, and minor wording suggestions for clarity). All retrieved references were independently verified by the authors, and all screening, interpretation, and citation decisions were made by the authors.

## Software availability

The analysis code is freely available at https://github.com/nathoogroup/SpaTran-NS. Per-gene results, dataset-level summaries, gene-set enrichment results, simulation summaries, and the software reproducibility manifest are provided as Supplemental files and described in Supplemental Table S7.

## Data access

The 12 publicly available datasets analyzed in this study are listed with repository, accession, and version information in Supplemental Table S2. No new sequencing data were generated in this study. The analysis code is available at https://github.com/nathoogroup/SpaTran-NS. Per-gene results, dataset-level summaries, gene-set enrichment results, simulation summaries, and the software reproducibility manifest are provided as Supplemental files and described in Supplemental Table S7.

## Competing interest statement

The authors declare no competing interests.

## Funding statement

Nathoo acknowledges funding from CIHR (PJT-186124), the Terry Fox Research Institute (TFRI1136-03), and NSERC (RGPIN-04044-2020). Velidi acknowledges funding from The Maud Menten Institute

## Acknowledgments

The authors acknowledge Céline Laumont and Brad H. Nelson for useful discussions related to spatial transcriptomic data, spatial aggregation of cells, and the results of the simulation studies.

## Author contributions

Conceptualization: FSN; Methodology: PV, ZW, FSN; Data analysis: PV; Data collection and preprocessing: PV; Software: PV; Supervision: FSN; Writing – original draft: PV, FSN; Writing – review and editing: PV, ZW, FSN.

## Notes

### Competing Interest Statement

The authors have declared no competing interest.

### Summary of Updates

Figures updated to not be blurry.

