## Supplementary Information for "Covariance Nonstationarity is Evident in Spatial Transcriptomics Providing a New Categorization of Spatially Varying Genes"

### S1 Abbreviations and Notation

Table S1: Abbreviations used in the supplementary material and additional files.

| Abbreviation | Expansion | Use |
|---|---|---|
| GO | Gene Ontology | Controlled vocabulary used for gene-set enrichment analysis. |
| GO ID | Gene Ontology identifier | Stable term identifier formatted as `GO:0000000`. |
| GO BP | Gene Ontology Biological Process | GO sub-ontology analyzed with `clusterProfiler`. |
| KEGG | Kyoto Encyclopedia of Genes and Genomes | Pathway database analyzed with `clusterProfiler` and MSigDB gene sets. |
| hsa / mmu | Human / mouse KEGG organism codes | KEGG organism prefixes used for *Homo sapiens* and *Mus musculus*. |
| GSEA | Gene set enrichment analysis | Rank-based enrichment test using genes ordered by $\log BF_{\mathrm{NS,S}}$. |
| ES | Enrichment score | Running-sum enrichment statistic reported by GSEA methods. |
| NES | Normalized enrichment score | ES normalized for gene-set size and permutation behavior. |
| FDR / padj | False discovery rate / adjusted $p$-value | Multiple-testing adjusted enrichment significance. |
| MSigDB | Molecular Signatures Database | Source of Hallmark, KEGG, Reactome, and GO gene-set collections used in the fgsea analysis. |
| fgsea | Fast gene set enrichment analysis | R package used for the MSigDB enrichment analysis. |
| INLA | Integrated Nested Laplace Approximation | Approximate Bayesian inference engine used for model fitting. |
| SPDE | Stochastic partial differential equation | Representation used to approximate Matern Gaussian random fields. |
| BF | Bayes factor | Ratio of model marginal likelihoods. |
| NS, S, NP | Nonstationary spatial, stationary spatial, stationary non-spatial | Model categories used in the Bayes factor comparisons. |

### S2 Supplementary Methods

#### S2.1 Datasets and quality control

Spatial transcriptomics datasets were obtained as SpatialExperiment objects and analyzed at the gene-by-spot count matrix level. Spots explicitly marked `in_tissue == 1` were retained; objects without an `in_tissue` field were treated as already tissue-restricted and retained all supplied spots. This removed 1,353 spots from `Visium_humanDLPFC`, leaving 3,639, and 2,290 spots from `Visium_mouseCoronal`, leaving 2,702. Re-release duplicates marked with `_v3.13` were excluded to avoid analyzing the same tissue sample twice. `MouseBrainCoronal` was also excluded in favor of the `Visium_mouseCoronal` representation of the same section. For each retained dataset, genes were kept when they were detected in at least 0.5% of analyzed tissue spots and had total count at least 3 across those spots. The resulting analysis set included 12 datasets spanning human and mouse tissues (Supplementary Table S2). Space Ranger versions were recorded from the 10x Genomics source datasets archived in ExperimentHub when available, or from the source documentation for the spatialLIBD DLPFC and STexampleData mouse coronal records.

#### S2.2 Spatial residual expression preprocessing and plotting

Spatial residual expression maps in the main manuscript and supplement show observed in-tissue Visium spots with detrended residuals below $-1$ or above $+1$. Spots with $|e_i| \leq 1$ are omitted from the display; they remain in the model fits. This fixed threshold is applied in residual units to every gene and dataset. The white plotting background is not an H&E image, tissue mask, or interpolated surface; blank areas can contain omitted spots or locations without measured spots and do not indicate zero expression.

For each representative gene, the plotted value at each spot was the same residual expression quantity used in the spatial covariance analysis: raw counts were transformed as

$$\log(1 + \text{count}),$$

rank-quantile normalized to normal scores, and detrended by subtracting the fitted quadratic coordinate trend $x + y + x^2 + y^2 + xy$. Points were plotted at spatial coordinates $(x, -y)$ with a fixed aspect ratio so that the plotted tissue orientation matched the image-like coordinate convention used by the input SpatialExperiment objects. Spots are shown without outlines; color encodes the residual value.

Colors represent actual detrended residual values on a shared diverging blue-white-red scale centered at zero, with fixed limits $[-2, 2]$. Negative residuals are blue (`#2166AC`), zero is white, and positive residuals are red (`#B2182B`). Residuals below $-2$ or above $+2$ remain displayed, with colors clipped to the corresponding endpoint. The shared numeric color bar is in normal-score residual units. White background denotes omitted or unmeasured spots, whereas white on the color bar denotes zero residual; zero-valued spots are excluded by the display threshold. Simulations use the same fixed threshold and color limits for their generated field values.

These maps show the locations and signs of residuals exceeding a common absolute threshold. The fraction of displayed spots can differ between genes and tissues. Model categories in Figures S2 and S3 are determined by the Bayes factor comparisons; visual clustering or an empty area alone does not establish covariance non-stationarity or identify a particular tissue compartment.

Figure 2 examples were selected within model-supported categories rather than solely by the largest Bayes factor. Non-stationary candidates required $BF_{\text{NS,S}} > 10$ and detection in at least 15% of tissue spots; up to 800 genes per dataset were screened in decreasing Bayes-factor order. For each spot, the 60 nearest neighbors defined local residual variance and the fraction with $|e| > 1$. Screening prioritized genes for which more than 8% of spots exceeded the residual threshold, more than 5% of neighborhoods had exceedance density below 0.05, and more than 5% had density above 0.30. Candidates were ranked by the product of these sparse- and dense-neighborhood fractions and $\log(1 + V_{90}/V_{10})$, where $V_{90}/V_{10}$ summarizes variation in local variance; final examples were chosen by inspecting maps. This diagnostic ratio does not set the display threshold or color limits. Non-spatial candidates required $BF_{\mathrm{S,NS}} > 10$ and $BF_{\mathrm{NP,S}} > 10$; up to 800 were screened in decreasing detection frequency. Among genes detected in at least 10% of spots, we evaluated the sum of absolute Moran's $I$ for residuals, absolute Moran's $I$ for squared residuals (both using six nearest neighbors with row-normalized weights), and the standard deviation of local exceedance density. All four selected non-spatial examples had $|I| < 0.051$ for both residual quantities. These are descriptive post-selection diagnostics, not additional hypothesis tests or proof of independence. Stationary-spatial examples were retained; shared mouse examples in Figure S3 match Figure 2, while other supplementary examples retain the original Bayes-factor-ranked selections. The gene manifest and screening diagnostics are provided with the plotting code.

The illustrative simulations in Fig. 2C were generated on a $31 \times 31$ regular grid using seed 24052026. The nonstationary realization used a Paciorek–Schervish-type Matérn covariance with smoothness $\nu = 1$,

$$C_{ij} = \sigma_i \sigma_j \frac{2\ell_i \ell_j}{\ell_i^2 + \ell_j^2} m_1\left(\frac{\sqrt{2}\,\delta_{ij}}{\sqrt{\ell_i^2 + \ell_j^2}}\right), \qquad m_1(r) = rK_1(r), \quad m_1(0) = 1.$$

At the vertical midline, the marginal standard deviation and correlation scale changed from $\sigma = 0.18$ and $\ell = 0.24$ on the left to $\sigma = 1.80$ and $\ell = 0.075$ on the right. The stationary comparator used the corresponding Matérn correlation with constant $\sigma = 1$ and $\ell = 0.13$; the independence comparator comprised independent standard Gaussian values. For each realization, only simulated values below $-1$ or above $+1$ were plotted. Colors encode the simulated field values on the same zero-centered scale clipped to $[-2, 2]$; values within $[-1, 1]$ were omitted.

##### S2.3 SPDE mesh construction

A single two-dimensional INLA mesh was built per dataset and reused for all genes in that dataset. Mesh construction used spot coordinates with inner and outer maximum edge lengths set to one fifth and one third of the larger tissue-coordinate range, respectively, and a cutoff set to one twentieth of the smaller coordinate range:

$$\texttt{max.edge} = \{\max(R_x, R_y)/5,\ \max(R_x, R_y)/3\}, \qquad \texttt{cutoff} = \min(R_x, R_y)/20,$$

where $R_x$ and $R_y$ are the ranges of the observed spot coordinates in the horizontal and vertical dimensions. Reusing the same mesh within a dataset ensured that per-gene Bayes factors compared models on the same spatial discretization. Dataset-specific mesh settings and mesh sizes are shown in Supplementary Table S3.

##### S2.4 Stationary and nonstationary Matern models

The stationary model used the SPDE representation of a Matern field with $\alpha = 2$, constant $\tau$, and constant $\kappa$:

$$\log \tau(s) = \theta_1, \qquad \log \kappa(s) = \theta_2.$$

The nonstationary model used the same likelihood and mesh, but allowed the spatial range parameter to vary linearly across scaled mesh coordinates:

$$\log \tau(s) = \theta_1, \qquad \log \kappa(s) = \theta_2 + \theta_3 \tilde{x}(s) + \theta_4 \tilde{y}(s).$$

Both models were fit with a Gaussian likelihood using R-INLA. For each fit, the marginal likelihood reported by INLA was retained along with posterior summaries of $\tau$, $\kappa$, spatial variance, residual variance, and practical range. For $\alpha = 2$ in two dimensions, the reported stationary-equivalent summaries use

$$\sigma_b^2 = \{4\pi\kappa^2\tau^2\}^{-1}, \qquad \rho = \sqrt{8}/\kappa.$$

#### S2.5 Model-fitting code excerpts

The complete implementation is provided in `hpc/analysis_functions.R`. The following code chunks show the core preprocessing, mesh construction, SPDE model definitions, INLA fitting call, and Bayes factor calculation used for each gene.

Listing 1: Preprocessing and shared SPDE mesh construction.

```
normalize_expression <- function(expr) {
  expr_rank <- rank(expr, ties.method = "average")
  qnorm((expr_rank - 0.5) / length(expr))
}

detrend_expression <- function(coords, expr) {
  x_c <- coords$x - mean(coords$x)
  y_c <- coords$y - mean(coords$y)
  fit <- lm(expr ~ x_c + y_c + I(x_c^2) + I(y_c^2) + I(x_c * y_c),
            data = data.frame(expr = expr, x_c = x_c, y_c = y_c))
  expr - predict(fit)
}

create_spde_mesh <- function(coords) {
  x_range <- diff(range(coords$x))
  y_range <- diff(range(coords$y))
  max_range <- max(x_range, y_range)
  min_range <- min(x_range, y_range)

  INLA::inla.mesh.2d(
    loc = as.matrix(coords[, c("x", "y")]),
    max.edge = c(max_range / 5, max_range / 3),
    cutoff = min_range / 20
  )
}

expr <- log1p(counts_mat[gene_idx, ])
expr_proc <- normalize_expression(expr)
expr_proc <- detrend_expression(coords, expr_proc)
mesh <- create_spde_mesh(coords)
```

Listing 2: Stationary Matern SPDE model and INLA fitting call.

```
fit_stationary_matern <- function(coords, expr, mesh) {
  spde <- INLA::inla.spde2.matern(
    mesh = mesh,
    alpha = 2,
    B.tau = matrix(c(0, 1, 0), nrow = 1, ncol = 3),
    B.kappa = matrix(c(0, 0, 1), nrow = 1, ncol = 3)
  )

  A <- INLA::inla.spde.make.A(
    mesh = mesh,
    loc = as.matrix(coords[, c("x", "y")])
  )
  s.index <- INLA::inla.spde.make.index(
    name = "spatial",
    n.spde = spde$n.spde
  )

  stack <- INLA::inla.stack(
```

```
    data = list(y = expr),
    A = list(A, 1),
    effects = list(s.index, list(intercept = rep(1, length(expr)))),
    tag = "est"
  )

  INLA::inla(
    y ~ -1 + intercept + f(spatial, model = spde),
    data = INLA::inla.stack.data(stack, spde = spde),
    family = "gaussian",
    control.predictor = list(A = INLA::inla.stack.A(stack), compute = TRUE),
    control.compute = list(config = TRUE, dic = TRUE, cpo = TRUE)
  )
}
```

Listing 3: Nonstationary Matern SPDE model and INLA fitting call.

```
fit_nonstationary_matern <- function(coords, expr, mesh) {
  n_mesh <- mesh$n
  mesh_x <- as.vector(scale(mesh$loc[, 1]))
  mesh_y <- as.vector(scale(mesh$loc[, 2]))

  B.tau <- matrix(c(0, 1, 0, 0, 0), nrow = n_mesh, ncol = 5, byrow = TRUE)
  B.kappa <- cbind(
    rep(0, n_mesh),
    rep(0, n_mesh),
    rep(1, n_mesh),
    mesh_x,
    mesh_y
  )

  spde <- INLA::inla.spde2.matern(
    mesh = mesh,
    alpha = 2,
    B.tau = B.tau,
    B.kappa = B.kappa
  )

  A <- INLA::inla.spde.make.A(
    mesh = mesh,
    loc = as.matrix(coords[, c("x", "y")])
  )
  s.index <- INLA::inla.spde.make.index(
    name = "spatial",
    n.spde = spde$n.spde
  )

  stack <- INLA::inla.stack(
    data = list(y = expr),
    A = list(A, 1),
    effects = list(s.index, list(intercept = rep(1, length(expr)))),
    tag = "est"
  )

  INLA::inla(
    y ~ -1 + intercept + f(spatial, model = spde),
    data = INLA::inla.stack.data(stack, spde = spde),
    family = "gaussian",
    control.predictor = list(A = INLA::inla.stack.A(stack), compute = TRUE),
    control.compute = list(config = TRUE, dic = TRUE, cpo = TRUE)
  )
}
```

Listing 4: Marginal likelihood Bayes factor and stationary-equivalent summaries.

```
fit_s <- fit_stationary_matern(coords, expr_proc, mesh)
fit_ns <- fit_nonstationary_matern(coords, expr_proc, mesh)

log_ml_stationary <- fit_s$mlik[1, 1]
```

```
log_ml_nonstationary <- fit_ns$mlik[1, 1]

log_bayes_factor <- log_ml_nonstationary - log_ml_stationary
bayes_factor <- exp(log_bayes_factor)

hyper <- fit_s$summary.hyperpar
theta1 <- hyper$mean[grep("Theta1 for spatial", rownames(hyper))]
theta2 <- hyper$mean[grep("Theta2 for spatial", rownames(hyper))]
tau <- exp(theta1)
kappa <- exp(theta2)

sigma_b_sq <- 1 / (4 * pi * kappa^2 * tau^2)
spatial_range <- sqrt(8) / kappa
```

#### S2.6 Bayes factor model comparisons

The primary comparison was nonstationary spatial covariance versus stationary spatial covariance:

$$\log BF_{\mathrm{NS,S}} = \log p(y \mid M_{\mathrm{NS}}) - \log p(y \mid M_{\mathrm{S}}).$$

Genes with $BF_{\mathrm{NS,S}} > 1$ were classified as favoring the nonstationary spatial model. Genes with $BF_{\mathrm{NS,S}} < 1$ were classified as favoring the stationary spatial model in the primary two-model comparison. Evidence strength was summarized using thresholds $BF = 3, 10, 30, 100$, corresponding to anecdotal, moderate, strong, very strong, and extreme evidence categories. For datasets with non-spatial fits available, genes favoring the stationary spatial model were further compared against a stationary non-spatial model by $\log BF_{\mathrm{S,NP}} = \log p(y \mid M_{\mathrm{S}}) - \log p(y \mid M_{\mathrm{NP}})$. This second comparison produced the final three-way categories: nonstationary spatial, stationary spatial, and stationary non-spatial. Genes for which an optimizer failed to converge were retained as explicit rows with missing model outputs and an error message. Dataset-level percentages and category summaries use successfully fitted genes as their denominator; Supplementary Table S2 reports failed-fit counts separately.

#### S2.7 Simulation design

Simulation studies were performed for Human Breast Cancer (ILC) and Human Ovarian Cancer to evaluate the false discovery rate and average power of Bayes factor thresholds. Successfully fitted, simulation-eligible genes with $BF_{\mathrm{NS,S}} \leq 1$ in the real-data analysis were treated as null genes and simulated from estimated parameters of the stationary Matern model from the real data. Similarly, genes with $BF_{\mathrm{NS,S}} > 1$ were treated as alternative genes and simulated from the fitted nonstationary SPDE model when fitted nonstationary coefficients were available.

Each simulation replicate generated expression for all simulation-eligible genes, refit the stationary and nonstationary models, and recomputed $BF_{\mathrm{NS,S}}$. One hundred replicates were aggregated per dataset. False discovery rate and average power were then evaluated across Bayes factor thresholds $c$, with the decision rule $BF_{\mathrm{NS,S}} > c$.

For dataset $d$, simulation replicate $r = 1, \ldots, R$, gene $g$, and Bayes factor threshold $c$, define the rejection rule

$$\delta_{gr}^{(d)}(c) = \mathbb{I}\left\{ BF_{gr}^{(d)} > c \right\}.$$

Let $\mathcal{H}_0^{(d)}$ and $\mathcal{H}_1^{(d)}$ denote the null and alternative genes in dataset $d$. The numbers of false positives and true positives in replicate $r$ are

$$FP_r^{(d)}(c) = \sum_{g \in \mathcal{H}_0^{(d)}} \delta_{gr}^{(d)}(c),$$

and

$$TP_r^{(d)}(c) = \sum_{g \in \mathcal{H}_1^{(d)}} \delta_{gr}^{(d)}(c).$$

The replicate-level false discovery proportion is

$$FDP_r^{(d)}(c) = \frac{FP_r^{(d)}(c)}{FP_r^{(d)}(c) + TP_r^{(d)}(c)},$$

with $FDP_r^{(d)}(c) = 0$ if $FP_r^{(d)}(c) + TP_r^{(d)}(c) = 0$.

The estimated false discovery rate is the average over simulation replicates:

$$\widehat{FDR}^{(d)}(c) = \frac{1}{R}\sum_{r=1}^{R} FDP_r^{(d)}(c).$$

The replicate-level power is

$$Power_r^{(d)}(c) = \frac{TP_r^{(d)}(c)}{|\mathcal{H}_1^{(d)}|}.$$

The estimated average power is

$$\widehat{Power}^{(d)}(c) = \frac{1}{R}\sum_{r=1}^{R} Power_r^{(d)}(c) = \frac{1}{R|\mathcal{H}_1^{(d)}|}\sum_{r=1}^{R}\sum_{g \in \mathcal{H}_1^{(d)}} \mathbb{I}\left\{BF_{gr}^{(d)} > c\right\}.$$

#### S2.8 Gene-set enrichment analysis

Gene-set enrichment analysis ranked genes by $\log BF_{\mathrm{NS,S}}$, so positive enrichment scores indicate gene sets enriched among genes favoring nonstationary covariance and negative enrichment scores indicate gene sets enriched among genes favoring stationary covariance. GO biological process and KEGG analyses were run with `clusterProfiler` using organism-specific annotation databases. A second enrichment analysis used MSigDB Hallmark, KEGG, Reactome, and GO biological process collections with `fgsea`. Full enrichment outputs are provided as Additional files 6 and 7.

### S3 Supplementary Tables

Table S2: Analyzed datasets, Bioconductor repository resources, ExperimentHub accessions, in-tissue spot selection, Space Ranger versions, and proportions of genes favoring the nonstationary spatial model or classified as spatially varying. Spatially varying includes nonstationary-spatial and stationary-spatial genes. Percentages use successfully fitted genes as the denominator; failed optimizer fits are retained as explicit missing-result rows and reported separately. EH9628 and EH9629 are the current replacements for the accessions used during analysis, EH9516 and EH9517, respectively.

| Dataset | Repository | Accession | Species | Tissue | Spots supplied | Spots analyzed | Off-tissue removed | Genes | Failed fits | Space Ranger | % NS | % spatial |
|---|---|---|---|---|---|---|---|---|---|---|---|---|
| HumanBreastCancerILC | ExperimentHub (TENxVisiumData) | EH6696 | Human | Breast cancer (ILC) | 4325 | 4325 | 0 | 16564 | 0 | 1.2.0 | 50.0 | 83.6 |
| Visium_humanDLPFC | ExperimentHub (STexampleData) | EH9628 | Human | DLPFC | 4992 | 3639 | 1353 | 15231 | 0 | 1.0.0 | 24.1 | 33.1 |
| HumanSpinalCord | ExperimentHub (TENxVisiumData) | EH6703 | Human | Spinal cord | 2812 | 2812 | 0 | 16057 | 1 | 1.2.0 | 23.6 | 42.6 |
| Visium_mouseCoronal | ExperimentHub (STexampleData) | EH9629 | Murine | Brain (coronal) | 4992 | 2702 | 2290 | 16982 | 52 | 1.0.0 | 19.0 | 68.5 |
| MouseBrainSagittalPosterior | ExperimentHub (TENxVisiumData) | EH6705 | Murine | Brain (posterior) | 6644 | 6644 | 0 | 16137 | 11 | 1.1.0 | 18.8 | 82.3 |
| MouseBrainSagittalAnterior | ExperimentHub (TENxVisiumData) | EH6706 | Murine | Brain (anterior) | 5520 | 5520 | 0 | 16431 | 28 | 1.1.0 | 14.7 | 76.8 |
| HumanColorectalCancer | ExperimentHub (TENxVisiumData) | EH6698 | Human | Colorectal cancer | 3138 | 3138 | 0 | 16236 | 1 | 1.2.0 | 13.7 | 73.2 |
| HumanGlioblastoma | ExperimentHub (TENxVisiumData) | EH6699 | Human | Glioblastoma | 3468 | 3468 | 0 | 17963 | 4 | 1.2.0 | 13.6 | 74.4 |
| HumanLymphNode | ExperimentHub (TENxVisiumData) | EH6701 | Human | Lymph node | 4035 | 4035 | 0 | 18295 | 5 | 1.1.0 | 12.4 | 47.9 |
| HumanOvarianCancer | ExperimentHub (TENxVisiumData) | EH6702 | Human | Ovarian cancer | 3493 | 3493 | 0 | 16684 | 0 | 1.2.0 | 5.3 | 57.1 |
| MouseKidneyCoronal | ExperimentHub (TENxVisiumData) | EH6707 | Murine | Kidney | 1438 | 1438 | 0 | 16343 | 0 | 1.1.0 | 5.0 | 41.1 |
| HumanHeart | ExperimentHub (TENxVisiumData) | EH6700 | Human | Heart | 4247 | 4247 | 0 | 13689 | 0 | 1.1.0 | 3.5 | 10.5 |

Table S3: INLA SPDE mesh settings used for each dataset. The mesh was generated with max.edge = c(max range / 5, max range / 3) and cutoff = min range / 20.

| Dataset | Spots | x range | y range | max.edge[1] | max.edge[2] | cutoff | Mesh nodes |
|---|---|---|---|---|---|---|---|
| HumanBreastCancerILC | 4325 | 18086 | 17081 | 3617.2 | 6028.7 | 854.0 | 313 |
| Visium_humanDLPFC | 3639 | 7966 | 9147 | 1829.4 | 3049.0 | 398.3 | 318 |
| HumanSpinalCord | 2812 | 2395 | 2639 | 527.8 | 879.7 | 119.8 | 360 |
| Visium_mouseCoronal | 2702 | 6605 | 8740 | 1748.0 | 2913.3 | 330.2 | 358 |
| MouseBrainSagittalPosterior | 6644 | 8640 | 8638 | 1728.0 | 2880.0 | 431.9 | 292 |
| MouseBrainSagittalAnterior | 5520 | 8130 | 8632 | 1726.4 | 2877.3 | 406.5 | 282 |
| HumanColorectalCancer | 3138 | 9234 | 7993 | 1846.8 | 3078.0 | 399.6 | 295 |
| HumanGlioblastoma | 3468 | 9231 | 7993 | 1846.2 | 3077.0 | 399.6 | 320 |
| HumanLymphNode | 4035 | 9239 | 8175 | 1847.8 | 3079.7 | 408.8 | 346 |
| HumanOvarianCancer | 3493 | 15768 | 17157 | 3431.4 | 5719.0 | 788.4 | 311 |
| MouseKidneyCoronal | 1438 | 6705 | 7992 | 1598.4 | 2664.0 | 335.2 | 221 |
| HumanHeart | 4247 | 8277 | 8618 | 1723.6 | 2872.7 | 413.9 | 320 |

Table S4: Simulation false discovery rate and average power at representative Bayes factor thresholds.

| Dataset | c | FDR | Power | Replicates | Genes | H0 genes | H1 genes |
|---|---|---|---|---|---|---|---|
| HumanBreastCancerILC | 1 | 0.003 | 0.874 | 100 | 16564 | 8284 | 8280 |
| HumanBreastCancerILC | 3 | 0.001 | 0.840 | 100 | 16564 | 8284 | 8280 |
| HumanBreastCancerILC | 10 | 0.001 | 0.798 | 100 | 16564 | 8284 | 8280 |
| HumanBreastCancerILC | 30 | 0.000 | 0.758 | 100 | 16564 | 8284 | 8280 |
| HumanBreastCancerILC | 100 | 0.000 | 0.712 | 100 | 16564 | 8284 | 8280 |
| HumanOvarianCancer | 1 | 0.081 | 0.811 | 100 | 16684 | 15803 | 881 |
| HumanOvarianCancer | 3 | 0.035 | 0.786 | 100 | 16684 | 15803 | 881 |
| HumanOvarianCancer | 10 | 0.016 | 0.757 | 100 | 16684 | 15803 | 881 |
| HumanOvarianCancer | 30 | 0.010 | 0.730 | 100 | 16684 | 15803 | 881 |
| HumanOvarianCancer | 100 | 0.007 | 0.700 | 100 | 16684 | 15803 | 881 |

Table S5: Top significant gene-set enrichment results from clusterProfiler.

| Dataset | Source | ID | Description | NES | FDR |
|---|---|---|---|---|---|
| HumanHeart | KEGG | hsa03010 | Ribosome | -4.057 | 0.000 |
| HumanHeart | KEGG | hsa00190 | Oxidative phosphorylation | -3.964 | 0.000 |
| HumanHeart | KEGG | hsa05012 | Parkinson disease | -3.419 | 0.000 |
| HumanHeart | KEGG | hsa05020 | Prion disease | -3.388 | 0.000 |
| HumanHeart | KEGG | hsa05171 | Coronavirus disease | -3.374 | 0.000 |
| HumanHeart | KEGG | hsa04260 | Cardiac muscle contraction | -3.304 | 0.000 |
| HumanHeart | KEGG | hsa05208 | Chemical carcinogenesis - reactive oxygen species | -3.206 | 0.000 |
| HumanHeart | KEGG | hsa04714 | Thermogenesis | -3.144 | 0.000 |
| HumanHeart | KEGG | hsa05415 | Diabetic cardiomyopathy | -3.119 | 0.000 |
| HumanHeart | KEGG | hsa04932 | Non-alcoholic fatty liver disease | -3.013 | 0.000 |
| HumanHeart | KEGG | hsa05016 | Huntington disease | -2.923 | 0.000 |
| HumanHeart | KEGG | hsa05014 | Amyotrophic lateral sclerosis | -2.685 | 0.000 |
| HumanHeart | KEGG | hsa05022 | Pathways of neurodegeneration - multiple diseases | -2.664 | 0.000 |
| HumanHeart | KEGG | hsa05010 | Alzheimer disease | -2.591 | 0.000 |
| HumanColorectalCancer | KEGG | hsa03040 | Spliceosome | -3.593 | 0.000 |
| HumanColorectalCancer | KEGG | hsa05012 | Parkinson disease | -3.431 | 0.000 |
| HumanColorectalCancer | KEGG | hsa00190 | Oxidative phosphorylation | -3.219 | 0.000 |
| HumanColorectalCancer | KEGG | hsa04640 | Hematopoietic cell lineage | 3.108 | 0.000 |
| HumanColorectalCancer | KEGG | hsa04061 | Viral protein interaction with cytokine and cytokine receptor | 3.052 | 0.000 |
| HumanColorectalCancer | KEGG | hsa04060 | Cytokine-cytokine receptor interaction | 3.026 | 0.000 |

Table S6: GO and KEGG identifier expansions for representative significant gene-set enrichment results. The complete identifier-to-description mapping is provided in Additional file 6.

| Source | Identifier | Full term or pathway name | Example dataset | NES | FDR |
|---|---|---|---|---|---|
| GO_BP | GO:0006402 | mRNA catabolic process | HumanColorectalCancer | -3.532 | 8.29e-09 |
| GO_BP | GO:0000956 | nuclear-transcribed mRNA catabolic process | HumanColorectalCancer | -3.439 | 8.29e-09 |
| GO_BP | GO:0042773 | ATP synthesis coupled electron transport | HumanColorectalCancer | -3.429 | 8.29e-09 |
| GO_BP | GO:0042775 | mitochondrial ATP synthesis coupled electron transport | HumanColorectalCancer | -3.429 | 8.29e-09 |
| GO_BP | GO:0000375 | RNA splicing, via transesterification reactions | HumanColorectalCancer | -3.378 | 8.29e-09 |
| GO_BP | GO:0022618 | protein-RNA complex assembly | HumanColorectalCancer | -3.372 | 8.29e-09 |
| GO_BP | GO:0071826 | protein-RNA complex organization | HumanColorectalCancer | -3.362 | 8.29e-09 |
| GO_BP | GO:0140053 | mitochondrial gene expression | HumanColorectalCancer | -3.325 | 8.29e-09 |
| GO_BP | GO:0019646 | aerobic electron transport chain | HumanColorectalCancer | -3.321 | 8.29e-09 |
| GO_BP | GO:0016072 | rRNA metabolic process | HumanColorectalCancer | -3.315 | 8.29e-09 |
| GO_BP | GO:0006119 | oxidative phosphorylation | HumanColorectalCancer | -3.284 | 8.29e-09 |
| GO_BP | GO:0000377 | RNA splicing, via transesterification reactions with bulged adenosine as nucleophile | HumanColorectalCancer | -3.252 | 8.29e-09 |
| KEGG | hsa03010 | Ribosome | HumanHeart | -4.057 | 2.36e-09 |
| KEGG | hsa00190 | Oxidative phosphorylation | HumanHeart | -3.964 | 2.36e-09 |
| KEGG | hsa05012 | Parkinson disease | HumanHeart | -3.419 | 2.36e-09 |
| KEGG | hsa05020 | Prion disease | HumanHeart | -3.388 | 2.36e-09 |
| KEGG | hsa05171 | Coronavirus disease | HumanHeart | -3.374 | 2.36e-09 |
| KEGG | hsa04260 | Cardiac muscle contraction | HumanHeart | -3.304 | 2.36e-09 |
| KEGG | hsa05208 | Chemical carcinogenesis - reactive oxygen species | HumanHeart | -3.206 | 2.36e-09 |
| KEGG | hsa04714 | Thermogenesis | HumanHeart | -3.144 | 2.36e-09 |
| KEGG | hsa05415 | Diabetic cardiomyopathy | HumanHeart | -3.119 | 2.36e-09 |
| KEGG | hsa04932 | Non-alcoholic fatty liver disease | HumanHeart | -3.013 | 2.36e-09 |
| KEGG | hsa05016 | Huntington disease | HumanHeart | -2.923 | 2.36e-09 |
| KEGG | hsa05014 | Amyotrophic lateral sclerosis | HumanHeart | -2.685 | 2.36e-09 |

Table S7: Additional files supplied with the manuscript.

| File | Format | Title | Description |
|---|---|---|---|
| Additional file 1 | PDF | Supplementary Information | Supplementary methods, figures, legends, and summary tables. |
| Additional file 2 | CSV | Dataset manifest and analysis summary | Dataset labels, repositories and accessions, supplied and analyzed in-tissue spot counts, removed off-tissue spot counts, numbers of genes and failed fits, and dataset-level model-category summaries. |
| Additional file 3 | CSV | Per-gene Bayes factor and category summary | Per-gene log Bayes factors, evidence strengths, nonstationary-versus-stationary model calls, available stationary-versus-non-spatial comparisons, and final model category. |
| Additional file 4 | CSV | Evidence-strength summary | Counts and percentages by dataset, favored model, and Bayes factor evidence strength for the nonstationary versus stationary comparison. |
| Additional file 5 | CSV | Stationary versus non-spatial summary | Counts and percentages of stationary-favored genes that further favor stationary spatial or stationary non-spatial models. |
| Additional file 6 | CSV | clusterProfiler GSEA results | GO biological process and KEGG enrichment results from genes ranked by $\log BF_{\mathrm{NS,S}}$. |
| Additional file 7 | CSV | MSigDB fgsea results | Hallmark, KEGG, Reactome, and GO biological process enrichment results from genes ranked by $\log BF_{\mathrm{NS,S}}$. |
| Additional file 8 | CSV | Simulation FDR and power curves | Aggregated false discovery rate and average power across Bayes factor thresholds. |
| Additional file 9 | CSV | Simulation replicate summaries | Per-replicate false discovery rate and power at $c = 1$. |
| Additional file 10 | CSV | Software reproducibility manifest | Analysis scripts, workflow roles, file sizes, and MD5 checksums. |

#### S4 Supplementary Figures

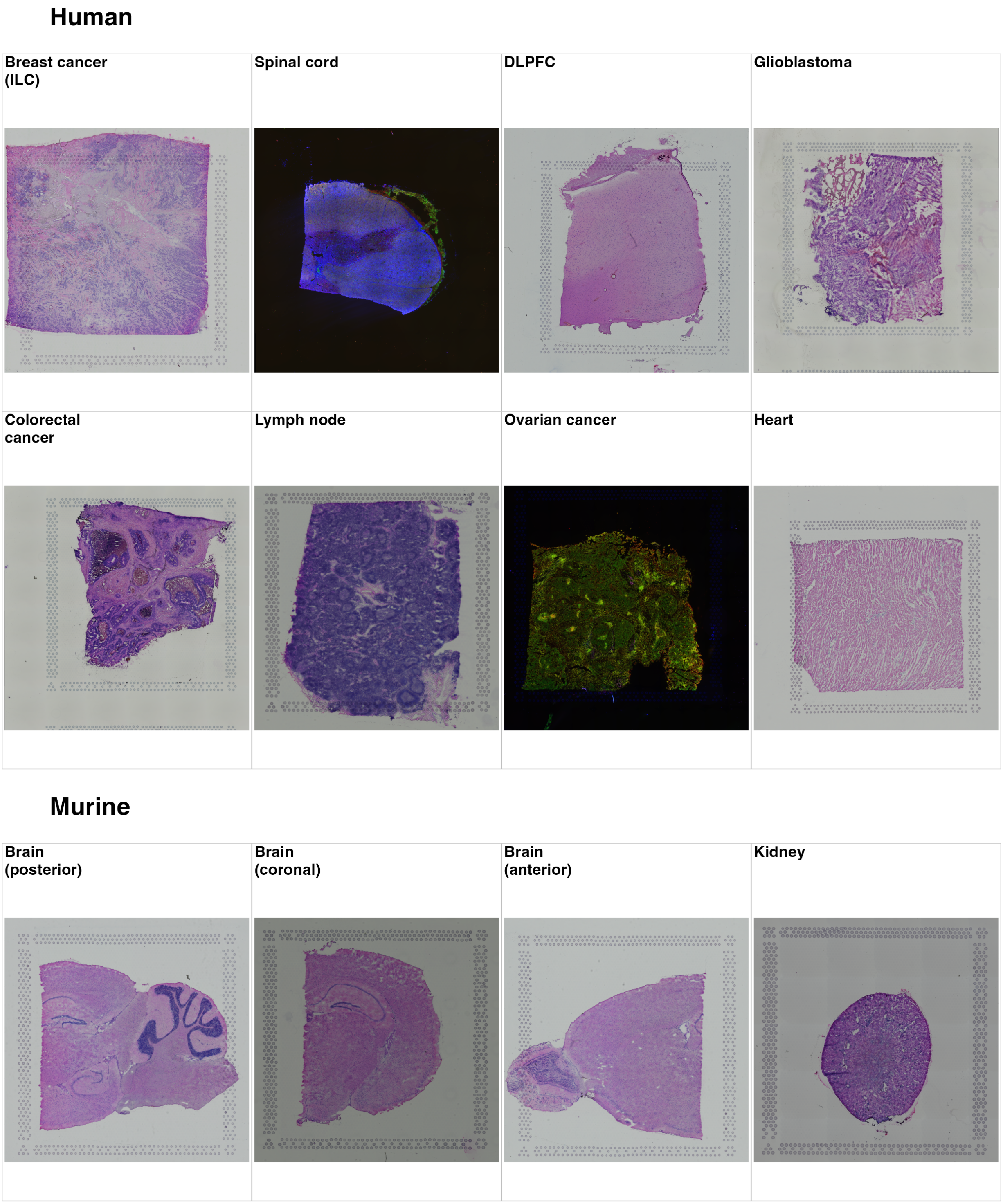


**Figure S1: Tissue image overview.** Low-resolution tissue images for the analyzed datasets. Images are the low-resolution tissue images stored in the corresponding ExperimentHub SpatialExperiment objects. Images are arranged by species and labeled by tissue/sample type.

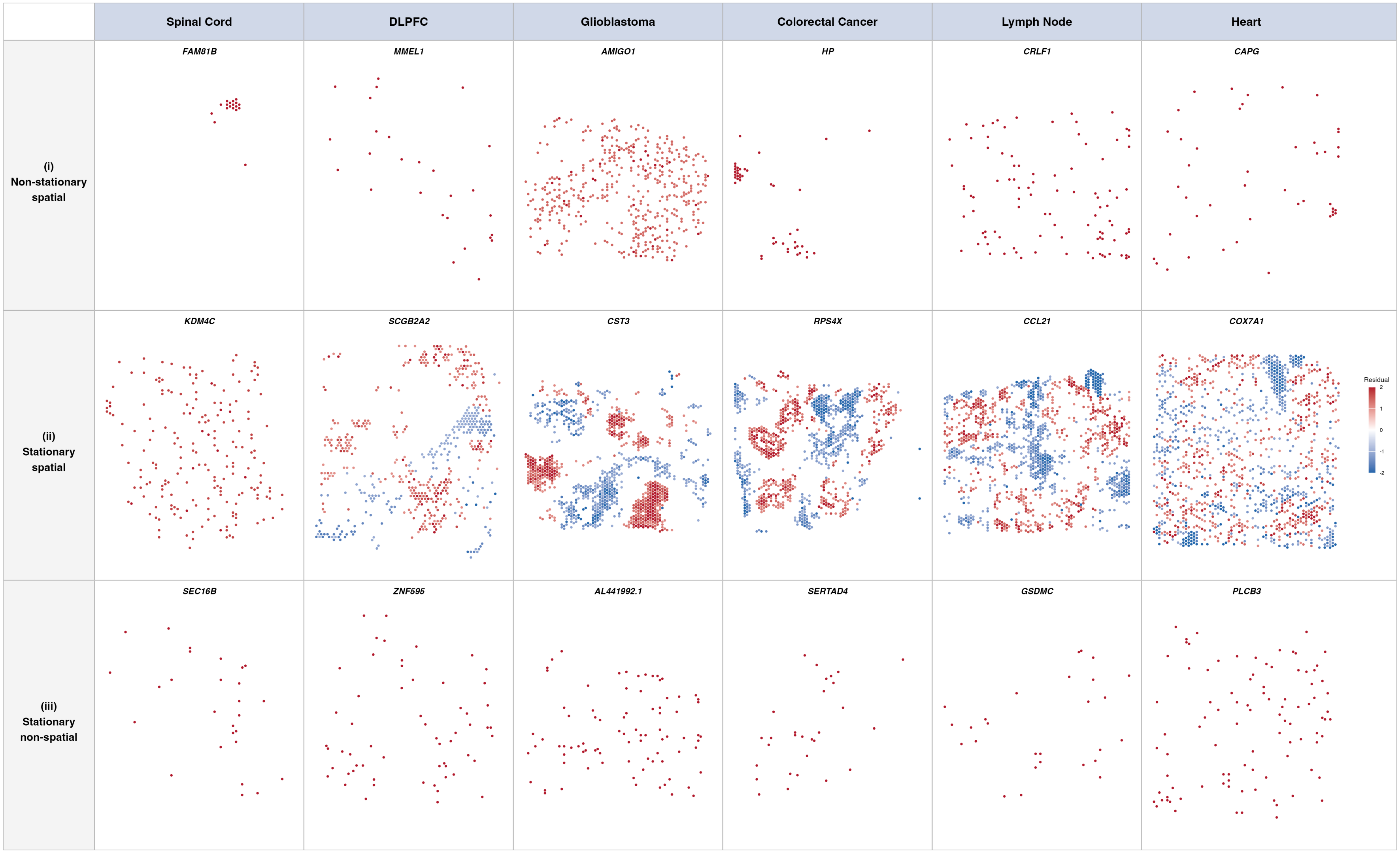


**Figure S2: Representative residual spatial maps for additional human datasets.** For each tissue, rows show representative genes from the nonstationary spatial, stationary spatial, and stationary non-spatial categories when available. Only residuals below $-1$ or above $+1$ are shown. Colors report actual residual values on the shared zero-centered scale clipped to $[-2, 2]$, matching the main manuscript. Values outside the color limits remain displayed at the endpoint colors.

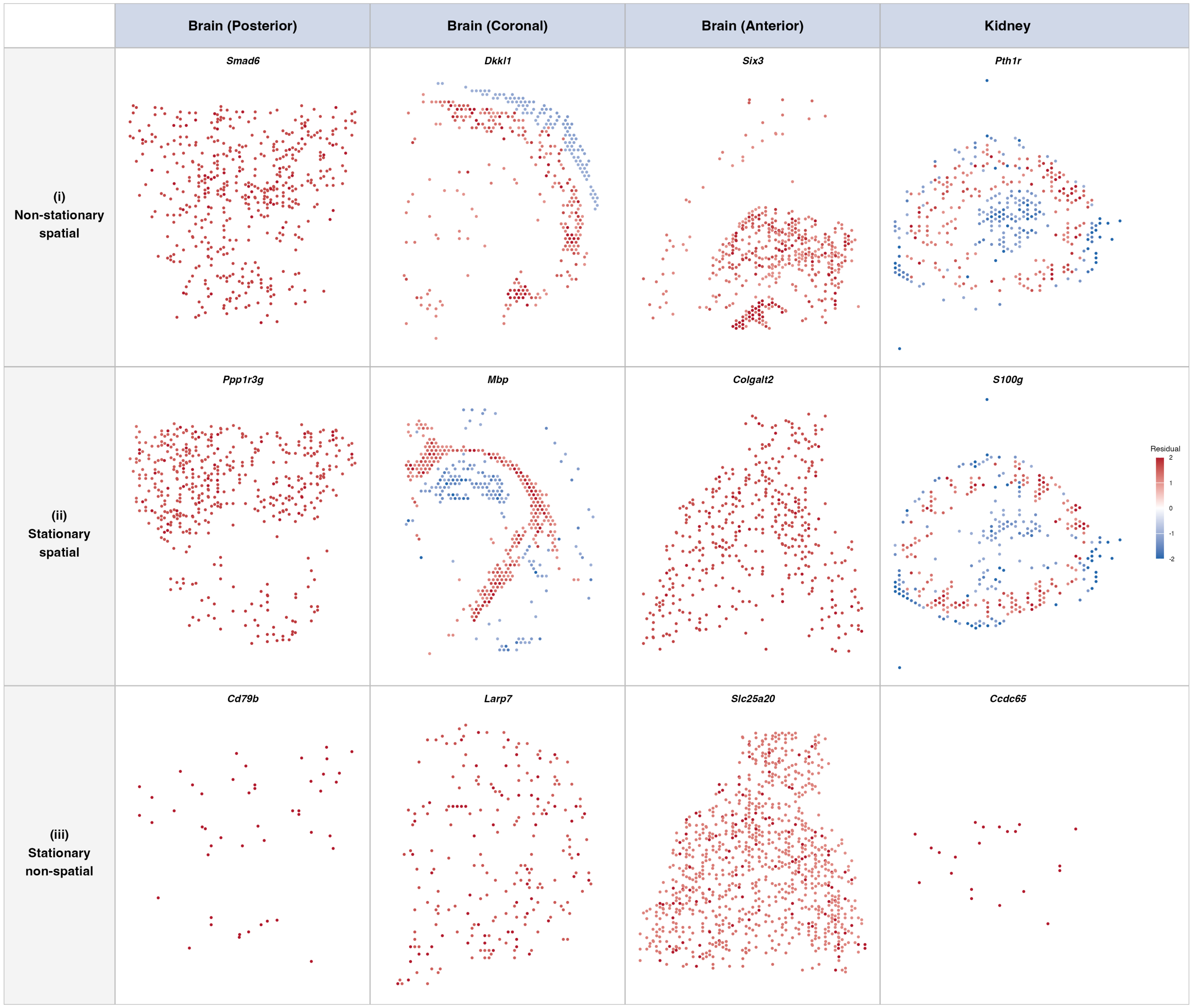


**Figure S3: Representative residual spatial maps for murine datasets.** For each tissue, rows show representative genes from the nonstationary spatial, stationary spatial, and stationary non-spatial categories when available. Only residuals below $-1$ or above $+1$ are shown. Colors report actual residual values on the shared zero-centered scale clipped to $[-2, 2]$, matching the main manuscript. Values outside the color limits remain displayed at the endpoint colors.

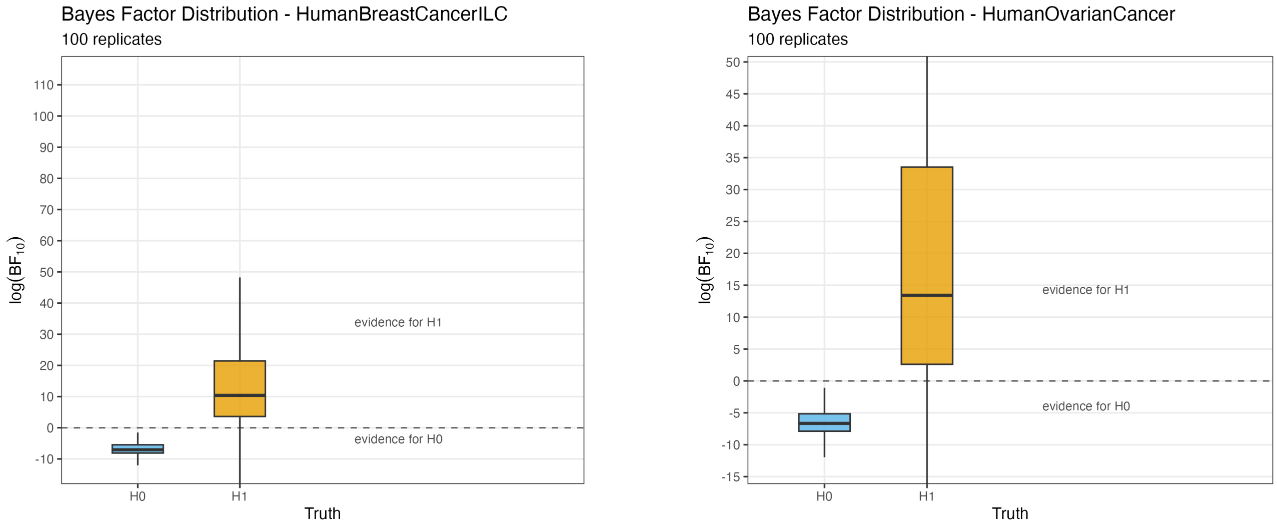


**Figure S4: Bayes factor distribution box plots in the two simulation source datasets.** Distributions show simulated log Bayes factors across 100 replicates each for Human Breast Cancer (ILC) and Human Ovarian Cancer. The $H_0$ and $H_1$ labels identify genes simulated from fitted stationary and nonstationary spatial models, respectively. Only finite Bayes-factor estimates are shown.
